# The genome of the coral model sea anemone *Exaiptasia diaphana* (Aiptasia) strain F003

**DOI:** 10.64898/2026.08.25.747183

**Authors:** Melanie Dörr, Abdoallah Sharaf, Luigi Colin, Kevin Schuster, Alyssa C. Bell, Christian R. Voolstra

## Abstract

We present a genome assembly of Aiptasia strain F003, a broadly used laboratory strain of the sea anemone and coral model organism *Exaiptasia diaphana* (Cnidaria; Anthozoa; Hexacorallia; Actiniaria; Aiptasiidae; *Exaiptasia*). The genome assembly spans 237.34 Mb across 12,480 contigs with a contig N50 of 76.47 kb (12,423 scaffolds with a scaffold N50 of 77.93 kb), including a single-contig mitochondrial genome with a length of 19.79 kb. The assembly is highly complete with a BUSCO completeness of 96.50% based on the metazoa dataset, including 94.80% single-copy, 1.70% duplicated, 1.70% fragmented, and 1.80% missing BUSCO genes. Genome annotation identified 29,589 protein-coding genes (including 2 pseudogenes) and a repeat content of 32.89%. The genome of the female Aiptasia strain F003 enhances the utility of a key cnidarian model organism by enabling comparisons among Aiptasia strains in studies of symbiosis, microbiomes, and thermal stress. It thereby strengthens the value of Aiptasia as a model for investigating the mechanisms underlying coral holobiont function, response, and resilience to environmental change.

## Introduction

*Exaiptasia diaphana*, commonly Aiptasia, (Eukaryota; Metazoa; Eumetazoa; Cnidaria; Anthozoa; Hexacorallia; Actiniaria; Aiptasiidae; *Exaiptasia*; NCBI:[txid2652724](https://www.ncbi.nlm.nih.gov/Taxonomy/Browser/wwwtax.cgi?id=2652724)) is a species of actiniarian sea anemones in the genus *Exaiptasia*, closely related to major reef-building stony corals (Scleractinia) within the class Hexacorallia in the cnidarian subphylum Anthozoa (Fig. 1A). Several *Exaiptasia diaphana* strains are used in laboratories worldwide (Fig. 1B), including H2 [1] originally obtained from Hawaii, F003 [2], and CC7 [3] originally collected in North Carolina, and strains from the Great Barrier Reef (GBR) [4], New Zealand [5], and the Red Sea [6].

**Figure 1.**
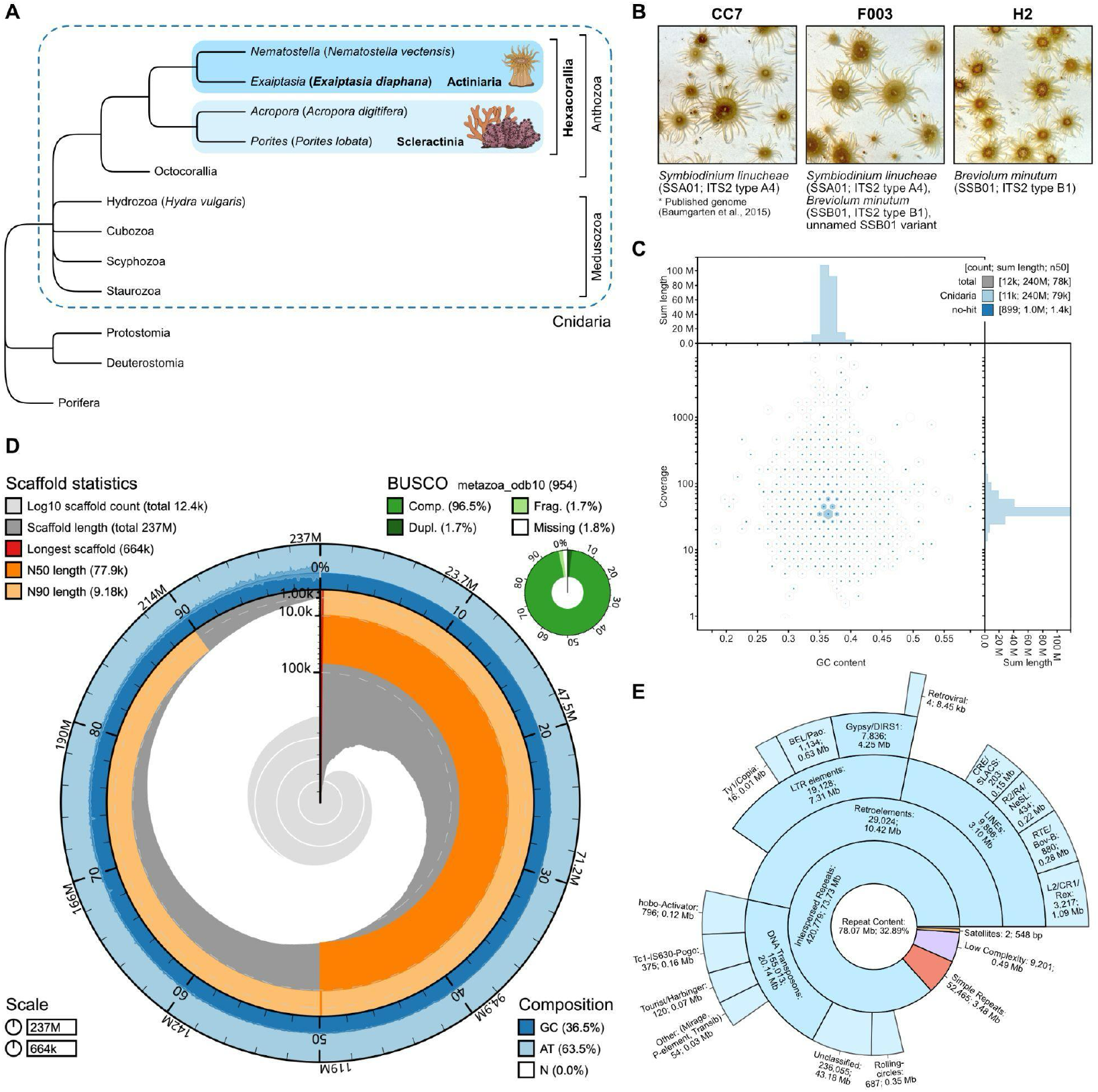
The genome of *Exaiptasia diaphana* (Aiptasia) strain F003. **(A)** Cladogram depicting the placement of *Exaiptasia diaphana* within the phylum Cnidaria. Actinitarian Aiptasia anemones are grouped together with sessile cnidarians, notably stony corals (Scleractinia), in the class of Hexacorallia. Representative species of various genera are noted in brackets. The cladogram is based on the NCBI taxonomy using Common Tree. The ggtree R package was used for visualization [36]. **(B)** Commonly used laboratory Aiptasia strains CC7, F003, and H2 with their respective algal symbiont compositions and ITS2 type majority sequences. Note that *Symbiodinium linucheae* is no longer considered valid under the International Code of Nomenclature (ICN) for Algae, Fungi, and Plants because the type specimen was a living culture rather than a permanent deposition [37]. Photo credits: Melanie Dörr. **(C)** GC-proportion square-binned blob plot (horizontal axis = GC content, vertical axis = sequence coverage) confirming the absence of putative contamination with contigs from other species. Blobs are sized proportionately to scaffold lengths and colored by phylum (light blue = Cnidaria, dark blue = no hit). **(D)** Genome assembly of Aiptasia strain F003. The BlobToolKit v4.1.2 Snailplot shows key assembly metrics and BUSCO gene completeness. The main plot is divided into 1,000 size-ordered bins around the circumference. Each bin represents 0.1% of the 237.34 Mb assembly. The central light grey spiral represents the cumulative scaffold count on a logarithmic scale. The dark grey spiral shows the distribution of scaffold lengths with the plot radius scaled to the longest scaffold (664 kb, red). The dark orange and light orange arcs indicate N50 (77.93 kb) and N90 (9.18 kb) scaffold lengths, respectively (also noted in the top left corner). The dark and light blue outer rings represent the GC (36.50%), AT (63.50%), and N (0%) composition (as displayed in the bottom right corner). The top right corner summarizes complete, fragmented, duplicated, and missing BUSCO genes (based on metazoa_odb10, v5.8.2, n = 954). The bottom left corner indicates the scale of the genome assembly (~237 Mb) and its longest scaffold (664 kb). **(E)** Repeat content of the Aiptasia strain F003 genome. A total of 78.07 Mb (32.89%) of the genome assembly consists of repeat elements. Unclassified repeats make up 43.18 Mb (18.19%) of the genome, while transposable elements make up 20.14 Mb (8.49%).

Just like reef-building corals, Aiptasia live in symbiosis with photosynthetic dinoflagellate algae (family Symbiodiniaceae) [7,8] and other microbial partners [9]. Algal symbiont associations vary between different Aiptasia strains, with strain F003 (female) hosting a mixed assemblage of three Symbiodiniaceae taxa, namely *Breviolum minutum* (SSB01, majority ITS2 type sequence B1), a thus far unnamed variant of SSB01, and *Symbiodinium linucheae* (SSA01, majority ITS2 type sequence A4), whereas H2 (female) and CC7 (male) anemones form single-symbiont associations with SSB01 and SSA01, respectively [1,2] (Fig. 1B).

The dinoflagellate endosymbiosis within hexacorallians, including reef-building stony corals and sea anemones, is vulnerable to rising sea temperatures and ocean acidification [10]. Coral bleaching, i.e., the loss of algal symbionts [11], is one of the main drivers of coral mortality and reef degradation [12]. Coral responses to environmental stress are also shaped by diverse other microbial partners that contribute to host physiology, nutrient cycling, and stress tolerance [13,14]. In particular, shifts in bacterial community composition have been recorded under thermal stress, and experimental microbiome manipulation has shown that beneficial bacteria can improve bleaching resilience and recovery in corals [15,16]. Yet, we lack a general understanding of the underlying molecular mechanisms [16]. This is, at least in part, due to the fact that coral research remains challenging: the endangered status of many coral species makes sampling prohibitive, long generation times complicate long-term (genetic) studies, and calcareous exoskeletons interfere with experimental manipulation [17].

To address these challenges, Aiptasia sea anemones have been used to study cnidarian-dinoflagellate symbiosis with published work dating back at least to the 1970s [7,18]. Unlike reef-building corals, which are colonial animals, Aiptasia are solitary, single-polyp sea anemones. They are (i) comparatively small, fast-growing, and easy to maintain under laboratory conditions, (ii) provide access to effectively unlimited numbers of clonal individuals, (iii) can be maintained in non-symbiotic (aposymbiotic or axenic) states (facultative symbiosis) [19], and (iv) are amenable to microscopic, molecular, and genetic manipulation [20,21]. Aiptasia thus enables simple maintenance, high-throughput experimentation under controlled laboratory conditions, and the disassembly and reassembly of defined holobiont configurations [19,22]. Consequently, Aiptasia sea anemones have grown into a powerful model poised to advance our understanding of the functional and mechanistic aspects underlying cnidarian-algal-microbiome interactions.

Recent genome sequencing efforts across Anthozoa continue to expand species-specific genomic resources for comparative and mechanistic studies of coral biology, including biomineralization, environmental adaptation, and resilience [23–25]. So far, the molecular resources available for the model Aiptasia include sequenced genomes and transcriptomes [26,27]. Yet, of the commonly used laboratory strains CC7, H2, and F003, only CC7 has a published genome [28]. To support the increasing use of Aiptasia strains F003 and H2 in conjunction with bacterial community analyses, thermal stress assays, and metagenomic sequencing [29,30], we provide the first genome sequence of *Exaiptasia diaphana* strain F003 to complement existing genomic resources and facilitate future holobiont research.

## Results and Discussion

### Genome Assembly and Repeat Elements Identification

The genome of *Exaiptasia diaphana* strain F003 was sequenced using one aposymbiotic adult polyp collected under long-term rearing conditions at the University of Konstanz. Genomic DNA was sequenced on an Oxford Nanopore Technologies (ONT) MinION platform using a FLO-MIN114.012 flow cell, generating 7,636,000 raw, unfiltered reads with a mean read length of 1,084.1 base pairs (bp) and a mean Phred-scaled quality score (Q score) of 15.4 (97.12% base-call accuracy). A total of 7,628,266 cleaned reads remained after adapter trimming with Porechop v0.2.4. Reads containing internal adapter sequences were treated as putative chimeras and discarded using the --discard_middle option. An initial 258 Mb assembly was decontaminated, and a quality assessment of the genome assembly confirmed a phylum-specific assembly and the effective removal of non-cnidarian sequences (Fig. 1C). Taxonomic classification of the removed contigs (n = 121) during the decontamination step showed that the majority were assigned to Pseudomonadota bacteria, with a smaller number assigned to Annelida, Mollusca, Echinodermata, Arthropoda, and Porifera (Data S1, Data S2). The final nuclear genome assembly spanned 237.34 Mb across 12,480 contigs and a GC content of 36.50% (Fig. 1D). The assembly size is consistent with and similar to the *Exaiptasia diaphana* strain CC7 genome, which spans 237 Mb [28]. This draft assembly represents version 1 (V1.0) for Aiptasia strain F003 and was generated from ONT long-read data alone. Although Hi-C data were not available for this study, chromosome-scale scaffolding represents a meaningful future upgrade toward a V2.0 assembly. The current assembly has a contig N50 of 76.47 kb. Although the dataset provided approximately 35x nominal coverage, the ONT reads had a read-length N50 of only 1.83 kb (maximum read length 428 kb). This relatively short read-length distribution likely limited the number of reads capable of spanning long or complex repetitive regions and was therefore probably a major constraint on assembly contiguity, consistent with prior benchmarking of ONT assemblies [31]. Despite the modest contig N50, the assembly retained high gene-space completeness (96.50% complete BUSCOs, see below), indicating that read length primarily constrained contiguity rather than overall genome recovery. Contigs were joined using Flye’s internal --scaffold function, which uses the assembly repeat graph together with the existing read alignments to connect contigs where the graph supports an unambiguous path across a gap. This procedure resulted in 12,423 scaffolds with a scaffold N50 of 77.93 kb. The longest scaffold spanned 664.23 kb (Table 1). The genome assembly is highly complete with a BUSCO completeness of 96.50% (single = 94.80%, duplicated = 1.70%, fragmented = 1.70%, and missing = 1.80% genes) based on the metazoa_odb10 reference dataset (Table 1, Fig. 1D). Repetitive sequences identified through a combined approach employing RepeatModeler and Extensive *de novo* TE Annotator (EDTA) comprised a total of 78.07 Mb (32.89%) of the assembly. The fraction is slightly higher than the fraction of repetitive sequences (26%) reported in the Aiptasia strain CC7 genome [28]. In the Aiptasia strain F003 assembly, unclassified repeats comprised 43.18 Mb (18.19%) of the genome, transposable elements made up 20.14 Mb (8.49%), and retroelements accounted for 10.42 Mb (4.39%) (Fig. 1E, Table S1).

**Table 1.**
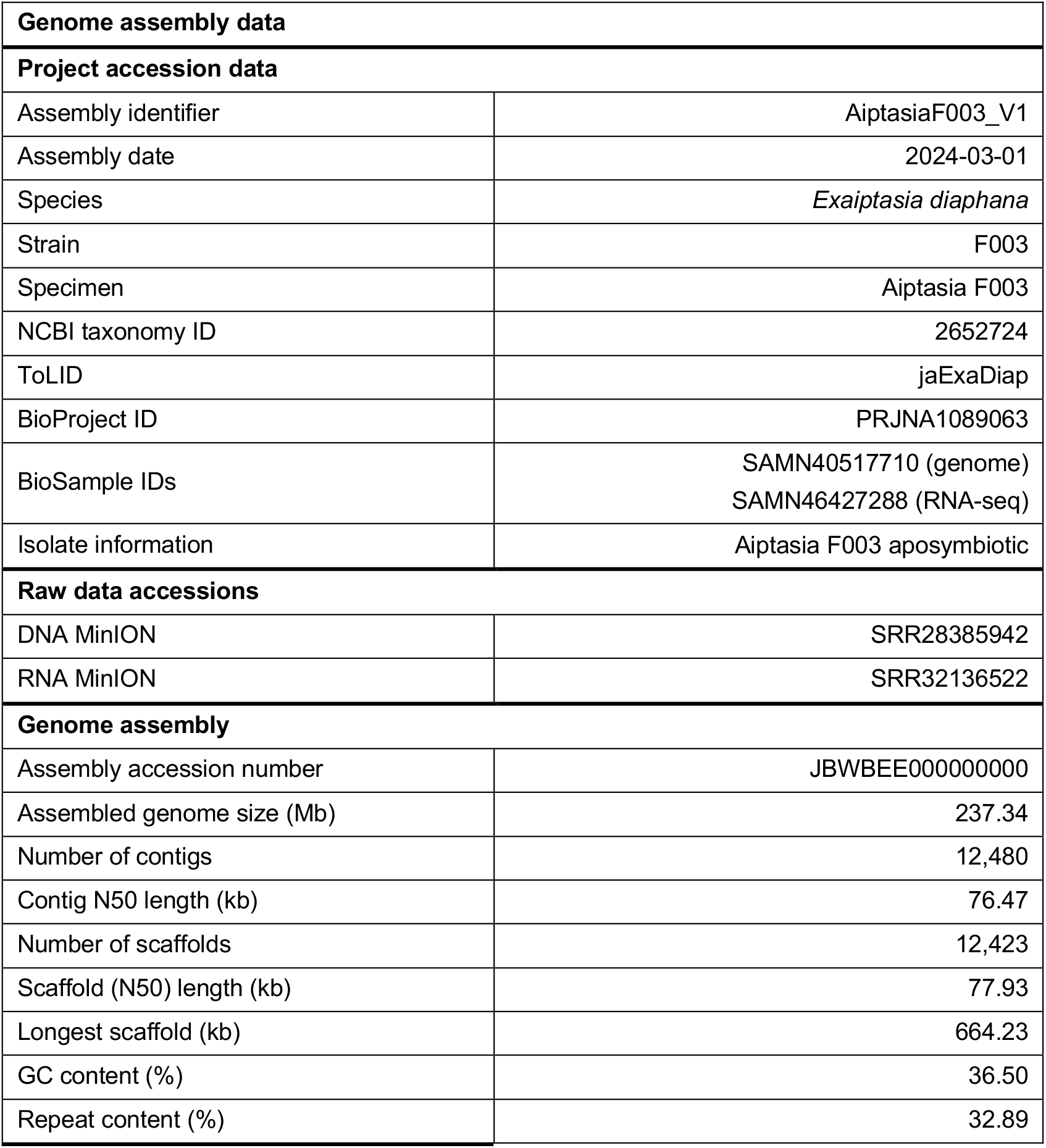
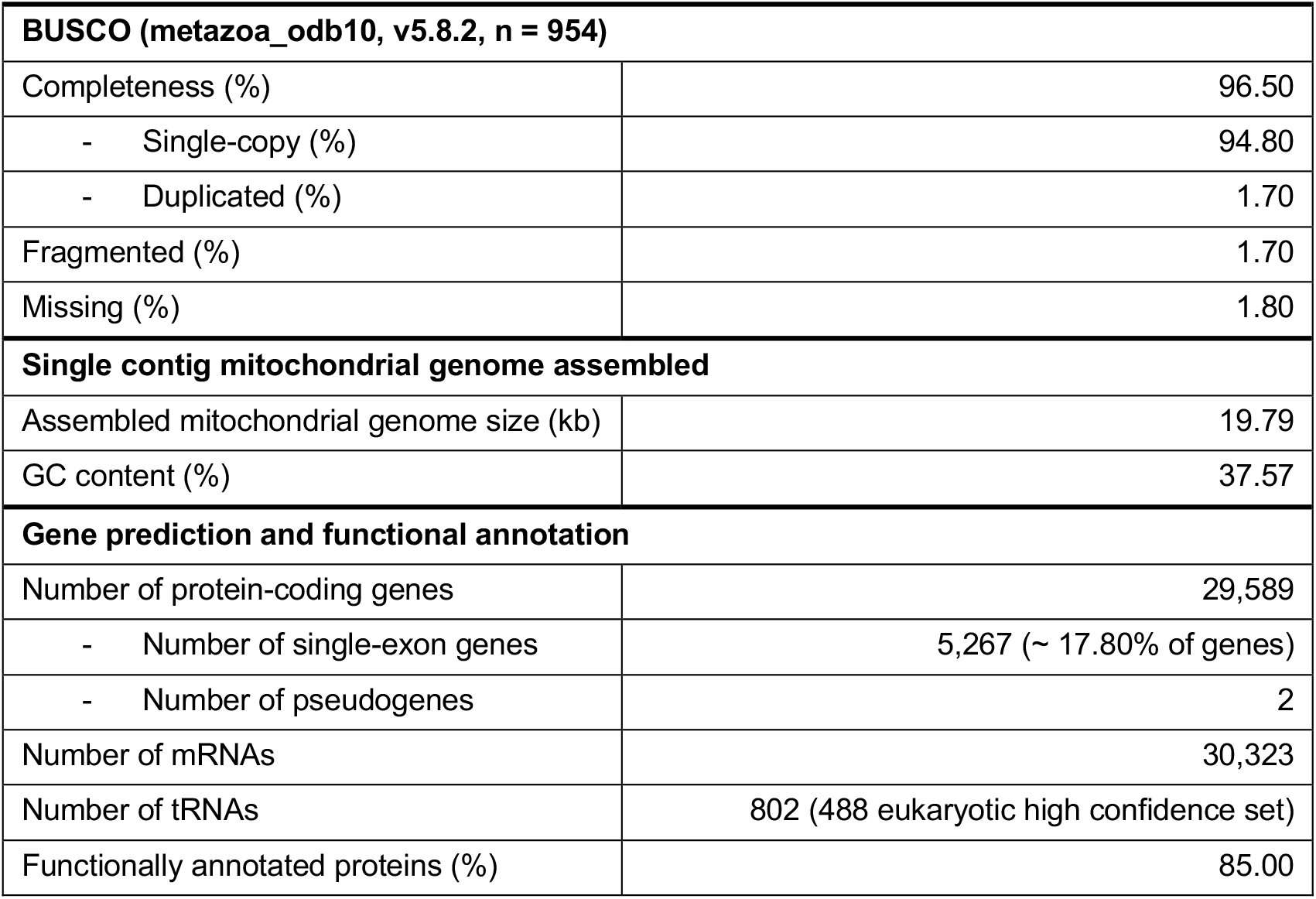
Genome assembly and functional annotation statistics of *Exaiptasia diaphana* (Aiptasia) strain F003. The percentage of functionally annotated proteins refers to the proportion of predicted proteins (aa) for which at least one functional annotation was successfully retrieved.

| Genome assembly data |  |
| --- | --- |
| Project accession data |  |
| Assembly identifier | AiptasiaF003_V1 |
| Assembly date | 2024-03-01 |
| Species | *Exaiptasia diaphana* |
| Strain | F003 |
| Specimen | Aiptasia F003 |
| NCBI taxonomy ID | 2652724 |
| ToLID | jaExaDiap |
| BioProject ID | PRJNA1089063 |
| BioSample IDs | SAMN40517710 (genome)<br>SAMN46427288 (RNA-seq) |
| Isolate information | Aiptasia F003 aposymbiotic |
| Raw data accessions |  |
| DNA MinION | SRR28385942 |
| RNA MinION | SRR32136522 |
| Genome assembly |  |
| Assembly accession number | JBWBEE000000000 |
| Assembled genome size (Mb) | 237.34 |
| Number of contigs | 12,480 |
| Contig N50 length (kb) | 76.47 |
| Number of scaffolds | 12,423 |
| Scaffold (N50) length (kb) | 77.93 |
| Longest scaffold (kb) | 664.23 |
| GC content (%) | 36.50 |
| Repeat content (%) | 32.89 |

| **BUSCO (metazoa_odb10, v5.8.2, n = 954)** |  |
| --- | --- |
| Completeness (%) | 96.50 |
| - Single-copy (%) | 94.80 |
| - Duplicated (%) | 1.70 |
| Fragmented (%) | 1.70 |
| Missing (%) | 1.80 |
| **Single contig mitochondrial genome assembled** |  |
| Assembled mitochondrial genome size (kb) | 19.79 |
| GC content (%) | 37.57 |
| **Gene prediction and functional annotation** |  |
| Number of protein-coding genes | 29,589 |
| - Number of single-exon genes | 5,267 (~ 17.80% of genes) |
| - Number of pseudogenes | 2 |
| Number of mRNAs | 30,323 |
| Number of tRNAs | 802 (488 eukaryotic high confidence set) |
| Functionally annotated proteins (%) | 85.00 |

## Mitogenome Assembly

The mitochondrial genome was assembled into a single contig of size 19.79 kb (Table 1), consistent with the circular mitochondrial genomes typically reported for actiniarian sea anemones [32]. Anthozoan actiniarian sea anemones are known for their variable mitogenome structure, with sizes varying from 16 kb (e.g., *Nematostella* sp.) to over 20 kb (e.g., *Urticina eques*). Interestingly, although it has been reported that GC content across nuclear loci in Hexacorallia is significantly higher than across mitochondrial loci [33], the Aiptasia mitogenome GC content (37.57%) was 1.07% higher compared to the nuclear genome assembly (36.50%) (Table 1). Other closely related taxa, such as the Zoantharia, have also been reported to exhibit slightly higher (0.5%) mitochondrial genome GC contents [33], and anthozoan mitogenomes are generally known to have higher GC content than medusozoans [34].

### Gene Prediction and Functional Annotation

To assist gene identification and annotation, total RNA of aposymbiotic F003 Aiptasia polyps was sequenced using the ONT MinION platform. A total of 29,589 protein-coding genes (including 2 pseudogenes) were identified, of which 27,152 (91.76%) represented complete gene models with start and stop codons (Table 1 and Table S2). Overall, 85.00% of the predicted protein-coding genes had functional annotations based on homology searches and domain assignments against multiple protein databases (Table 1). Here, the annotation rate refers to the proportion of predicted proteins for which at least one functional annotation was successfully retrieved. Functional annotation was assigned by comparing translated protein sequences to known protein families, conserved domains, and orthologous groups using tools such as InterProScan, eggNOG-mapper, and COGs (Table S2). As for the genome size, the number of genes is very similar to the *Exaiptasia diaphana* strain CC7 genome, which totals 29,269 genes [28]. Comparative orthology analysis using OrthoVenn3 [35] identified 16,548 orthologous groups (OGs), corresponding to 17,867 shared protein sequences between F003 and CC7, including 10,493 single-copy orthologs. F003 contained more strain-specific OGs (822) than CC7 (497), suggesting greater gene repertoire expansion in F003 (Figure S1). The F003 strain-specific gene models identified through the OrthoVenn3 orthology analysis together with their orthogroup assignments and available functional annotations are provided in the supplement (Data S3).

## Materials and Methods

### Aiptasia rearing and bleaching

Symbiotic anemones were reared in artificial seawater (ASW; PRO-REEF Sea Salt, Tropic Marin) at 35 gL^-1^ salinity and 25 °C with a 12-hour light, 12-hour dark cycle (~70-80 µmol photons m^-2^s^-1^ light intensity). Animals were fed once per week with freshly hatched *Artemia* nauplii (Ocean Nutrition). One day after feeding, anemone tanks were cleaned, and rearing water was exchanged. Aposymbiotic Aiptasia polyps of the clonal strain F003 were generated using a menthol/diuron treatment as previously described [38]. Briefly, menthol (20% w/v in ethanol) was added to 1 µm-filtered ASW to a final concentration of 0.19 mmol L^-1^. The anemones were incubated in the menthol/ASW solution for 8 hours during the 12-hour light period. For the following 16 hours, the anemones were incubated in a fresh solution of 1 µm-filtered ASW containing diuron (DCMU) (100 mM in ethanol) at a final concentration of 5 µM L^-1^ to inhibit the re-establishment of the algal symbiosis. The 24-hour treatment was repeated for four consecutive days, followed by a three-day break during which the anemones were kept in fresh ASW. During the seven-day treatment, the anemones were maintained under rearing conditions. The seven-day treatment cycle was repeated twice, after which the anemones were kept in darkness until sampling and subsequent nucleic acid extraction. Successful bleaching, i.e., expulsion of Symbiodiniaceae, was confirmed using a Zeiss Stemi 2000-C stereomicroscope equipped with a fluorescence green longpass (LP) filter adapter (excitation 510-540 nm, emission 600 nm LP) and UV light illuminator (NIGHTSEA).

### DNA extraction and sequencing

DNA extraction was conducted at the Sequencing Analysis Core Facility (SequAna) at the University of Konstanz. One aposymbiotic Aiptasia anemone of the clonal strain F003 was rinsed in 1X PBS and homogenized using a Polytron PT 1200 E homogenizer (Kinematica, Switzerland). DNA was extracted using the DNeasy Blood & Tissue Kit (Qiagen, Hilden, Germany) according to the manufacturer’s instructions. The Qubit dsDNA High Sensitivity Assay Kit (Thermo Fisher Scientific, Waltham, Massachusetts, USA) was used to assess DNA quantity. The DNA library was prepared with 300 fmol of gDNA following the manufacturer’s protocol for the Ligation Sequencing Kit V14 (SQK - LSK114, Oxford Nanopore Technologies, Oxford, UK). DNA sequencing was performed using the Oxford Nanopore Technologies (ONT) MinION Mk1B platform (Oxford Nanopore Technologies, Oxford, UK) with a FLO-MIN114.012 flow cell and super-accurate base-calling (dna_r10.4.1_e8.2_400bps_sup@v4.2.0). The flow cell was run for 72 h following the standard manufacturer’s protocol.

### RNA extraction and sequencing

RNA extraction was conducted at the Sequencing Analysis Core Facility (SequAna) at the University of Konstanz. Total RNA was extracted from 2 pools of 5 aposymbiotic F003 Aiptasia anemones using the RNeasy Mini Kit (Qiagen, Hilden, Germany). First, animals were lysed and homogenized in 600 µL Buffer RLT using a Polytron PT 1200 E homogenizer (Kinematica, Switzerland). Next, total RNA was extracted with an additional on-column DNAse I digestion (RNase-Free DNase Set, Qiagen, Hilden, Germany) according to the manufacturer’s instructions. Both RNA extractions were concentrated via lithium chloride precipitation. Briefly, an equal volume of 7.5 M lithium chloride was added to each sample, and the samples were incubated overnight at −20 °C. After centrifugation (12,000 x g, 15 min), the supernatant was removed. The pellets were washed with 2.5 sample volumes of 70% ethanol, centrifuged (12,000 x g, 2 min), air-dried (30 min), and dissolved in RNase-free water. RNA quantity and quality were assessed using the Qubit RNA High Sensitivity Assay Kit (Thermo Fisher Scientific, Waltham, Massachusetts, USA), NanoDrop spectrophotometry, and 1% agarose gel electrophoresis. The two RNA extractions were pooled into a single sample, and a total RNA-Seq library was constructed using the Direct RNA Sequencing Kit (SQK-RNA004, Oxford Nanopore Technologies, Oxford, UK) according to the manufacturer’s protocol. Direct RNA sequencing was performed on the ONT MinION platform, using a FLO-MIN004RA flow cell and super-accurate base-calling (rna004_130bps_sup@v3.0.1). As for DNA sequencing, the flow cell was run for 72 h following the standard manufacturer’s protocol.

### Nuclear and mitochondrial genome assembly and evaluation

Adapters were trimmed from the ONT DNA sequencing reads using Porechop v0.2.4 (RRID:SCR_016967), and reads were assembled using NECAT v0.0.1 (RRID:SCR_025350) [39], CANU v2.2 (RRID:SCR_015880) [40], and Flye v2.9.3 (RRID:SCR_017016) [41] (Table S3). NECAT and Canu incorporate read-correction procedures into their assembly workflows through progressive two-step correction and adaptive k-mer-weighted overlap correction, respectively. In contrast, Flye constructs a repeat graph directly from uncorrected reads and performs assembly consensus polishing of the resulting assembly using read alignments. Flye was run with --genome-size 275m and --scaffold; all other parameters were left at their default settings. The Flye assembly was chosen for downstream analysis since it showed the best assembly statistics [42]. Genome assembly characteristics were evaluated and visualized as taxon-annotated Guanine-Cytosine (GC)-proportion plots using BlobToolKit v4.1.2 (RRID:SCR_025882) [43]. The putative taxonomic identity of contigs was determined with searches against the National Center for Biotechnology Information (NCBI) non-redundant nucleotide database using BLASTn v2.14.1 (RRID:SCR_001598) and the UniProt database with DIAMOND v2.1.8 (RRID:SCR_016071) [44], following the BlobToolKit instructions. The initial assembly was decontaminated by removing non-cnidarian contigs based on GC content, coverage, and taxonomic assignment information provided by BlobToolKit. The completeness and contiguity of the assembly were evaluated using BUSCO v5.8.2 (RRID:SCR_015008) [45], whereby the Archaea and Bacteria datasets were used to assess decontamination and the Eukaryota and Metazoa OrthoDB v10 datasets were used to assess completeness across a broad evolutionary range. The mitochondrial genome was assembled using GetOrganelle v1.7.7.1 (RRID:SCR_022963) [46], while MITOS v2.1.0 [47] was used to obtain functional annotation of the mitogenomic sequence.

### DNA repeat identification

Genomic DNA repeats were identified using a combination of RepeatModeler v2.0.4 (RRID:SCR_015027) [48] and the Extensive *de novo* TE Annotator (EDTA) v2.2.0 (RRID:SCR_022063) [49] (Table S3). Identified repeats were masked with RepeatMasker v4.1.6 (RRID:SCR_012954) [50] using custom *Exaiptasia*-specific repeat libraries under the “LTRStruct” option, while default settings were applied for all other parameters. To visualize the hierarchical distribution of repeat elements, we utilized a sunburst plot using the Plotly Express library in Python (Plotly Technologies Inc.).

### Gene prediction and functional annotation

All gene prediction and functional annotation steps were orchestrated using our fully automated pipeline, GeneForge v1.0 [51], which supports both BRAKER v3.0.8 (RRID:SCR_018964) [52] and funannotate v1.8.15 (RRID:SCR_023039) [53]. GeneForge evaluated the completeness of the gene set annotations generated by BRAKER and funannotate using BUSCO, based on the corresponding predicted protein sequences. The gene models predicted using funannotate achieved the highest BUSCO completeness and were therefore selected for all downstream analyses (Table S3). As part of funannotate v1.8.15 (RRID:SCR_023039), five ab initio predictors were run independently: AUGUSTUS (RRID:SCR_008417), SNAP (RRID:SCR_007936), glimmerHMM (RRID:SCR_002654), CodingQuarry, and GeneMark-ES/ET [54–59]. Their outputs were then compiled into a single consensus gene set using EVidenceModeler (EVM) (RRID:SCR_014659) [60], which weights each ab initio prediction against the aligned RNA-seq evidence (see below) to resolve conflicting gene models. RNA-seq evidence came from two sources and served distinct roles. ONT direct RNA sequencing of aposymbiotic F003 polyps generated in this study was trimmed using Porechop v0.2.4 (RRID:SCR_016967) (Table 3), aligned to the assembled genome using STAR v2.7.11b (RRID:SCR_004463) [61], and assembled with StringTie v2.2.1 (RRID:SCR_016323) [62]. The resulting long-read transcript models were used to extend UTR boundaries and support splice sites during EVM weighting. Published Illumina RNA-seq data from aposymbiotic Aiptasia larval cells [63] provided complementary, independent short-read transcript evidence for gene models with limited or no support from the ONT dataset. All predicted proteins were functionally annotated using a combination of databases and tools. General functional annotation was performed with InterProScan v5.72-103.0 (RRID:SCR_005829) [64], eggNOG-mapper v2.1.12 (RRID:SCR_021165) [65,66], BUSCO v5.8.2 (RRID:SCR_015008) [45], Pfam (RRID:SCR_004726) [67], and COGs (RRID:SCR_007139). Carbohydrate-active enzyme families were identified using the CAZy database (RRID:SCR_012909) [68], while proteases were classified using MEROPS (RRID:SCR_007777). Moreover, we searched NCBI and UniProtKB/SwissProt databases using BLASTp v2.14.1 (RRID:SCR_001010) and DIAMOND v2.1.8 (RRID:SCR_016071) software for gene names and gene product descriptions [44,69]. Lastly, we predicted transmembrane topology and signal peptides using Phobius v1.01 (RRID:SCR_015643) and SignalP v6.0 (RRID:SCR_015644) [70,71].

## Availability of source code

All bioinformatic tools used in this study are listed in Table S3; all curated pipelines are available on GitHub at [https://github.com/SequAna-Ukon/Aiptasia_F003_genome] (MIT license). The pipelines are implemented in Bash and run on Linux operating systems.

## Data availability

Sequencing data and genome assembly (genome assembly ID: GCA_056151815.1; https://www.ncbi.nlm.nih.gov/datasets/genome/GCA_056151815.1/) are available under NCBI BioProject ID PRJNA1089063 (https://www.ncbi.nlm.nih.gov/bioproject/PRJNA1089063). The genome assembly FASTA file, coding gene annotation GFF file, coding gene nucleotide and protein sequence FASTA files, repeat/transposable element annotation GFF file, and BUSCO output files are available through Zenodo at https://doi.org/10.5281/zenodo.21918813 [72].

## Supplementary Material

Figure S1, Table S1, Table S2, Table S3, Data S1, Data S2, Data S3

## Ethics approval and consent for publication

Not applicable.

## Competing interests

No competing interests were disclosed.

## Authors’ contributions

MD, AS, CRV analyzed data; AS assembled and annotated the genome; LC, KS, ACB processed samples and generated sequencing data; CRV provided tools, reagents, funding. MD with AS, CRV wrote the manuscript with contributions from all authors.

## Funding

CRV acknowledges AFF funding by the University of Konstanz (Project INTEGER; grant number 15902919).

## Acknowledgments

We acknowledge the SequAna Core Facility BIO-16840 Genomics Practical Course for providing the platform to generate the data used in this study. AS is supported by the Department of Biology, University of Konstanz.

## Notes

### Competing Interest Statement

The authors have declared no competing interest.

