## Supplementary Material for "The genome of the coral model sea anemone *Exaiptasia diaphana* (Aiptasia) strain F003"

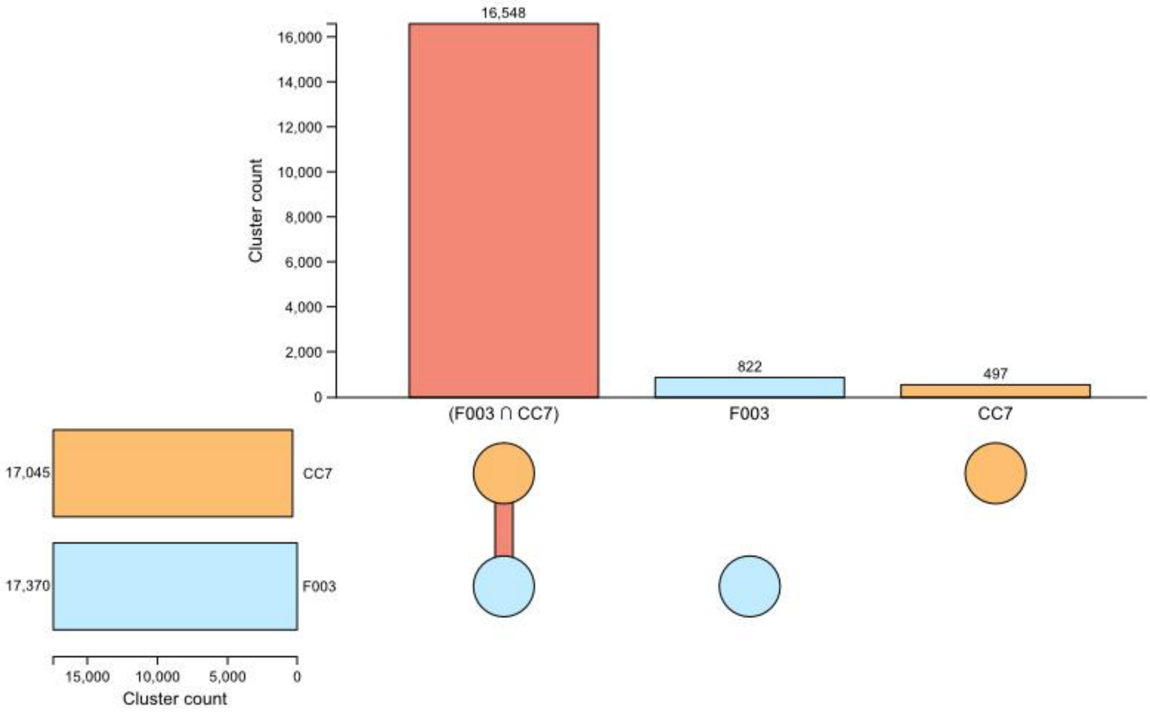


**Figure S1. Comparative orthology analysis of *Exaiptasia diaphana* (Aiptasia) strain F003 and CC7 genomes.** OrthoVenn3 identified 16,548 shared orthologous groups (OGs) (red). F003 contained more strain-specific OGs (light blue) than CC7 (light orange), indicating differences in the predicted gene repertoires of the two strains.

**Table S1. Repeat content of the *Exaiptasia diaphana* (Aiptasia) strain F003 genome assembly.** A total of 78.07 Mb (32.89%) of the genome assembly consists of repetitive sequences.

| **Repetitive content** | | |
|---|---|---|
| **Group, repeat type, and subtype** | **Number of elements** | **Length occupied (Mb)** |
| Interspersed repeats | 420,779 | 73.73 |
| DNA transposons | 155,013 | 20.14 |
| - Tc1-IS630-Pogo | 375 | 0.16 |
| - hobo-Activator | 796 | 0.12 |
| - Tourist/Harbinger | 120 | 0.07 |
| - Other (Mirage, P-element, Transib) | 54 | 0.03 |
| Retroelements | 29,024 | 10.42 |
| - LTR elements | 19,128 | 7.31 |
| - Gypsy/DIRS1 | 7,836 | 4.25 |
| - BEL/Pao | 1,134 | 0.63 |
| - Ty1/Copia | 16 | 0.01 |
| - Retroviral | 4 | 0.008 |
| - LINEs | 9,896 | 3.10 |
| - L2/CR1/Rex | 3,217 | 1.09 |
| - RTE/Bov-B | 880 | 0.28 |
| - R2/R4/NeSL | 434 | 0.22 |
| - CRE/SLACS | 203 | 0.15 |
| Simple repeats | 52,465 | 3.48 |
| Low complexity | 9,201 | 0.49 |
| Rolling-circles | 687 | 0.35 |
| Satellites | 2 | 0.0005 |
| Unclassified | 236,055 | 43.18 |

**Table S2. Additional gene prediction and functional annotation statistics of the *Exaiptasia diaphana* (Aiptasia) strain F003 genome assembly.** Summary of general annotation statistics, including functional annotations, as well as features of coding sequences (CDS), exons, introns, and untranslated regions (UTRs).

| **Gene prediction and functional annotation** | |
|---|---|
| Number of protein-coding genes | 29,589 |
| Number of mRNAs | 30,323 |
| **UTR statistics** | |
| Number of mRNAs with UTRs on both sides | 7,411 (~ 24.44% of mRNAs) |
| Number of mRNAs with at least one UTR | 11,249 (~ 37.09% of mRNAs) |
| Number of 5’ UTRs | 11,156 |
| Number of 3’ UTRs | 9,528 |
| **Exon and intron features** | |
| Total number of exons | 192,046 |
| Mean number of exons per gene | 6.30 |
| Total number of introns in CDS | 159,653 |
| Mean number of introns per CDS | 5.30 |
| **Lengths** | |
| Longest gene (bp) | 66,488 |
| Mean gene length (bp) | 3,617 |
| Longest mRNA (bp) | 66,488 |
| Longest CDS (bp) | 57,912 |
| Mean CDS length (bp) | 1,302 |
| Mean exon length (bp) | 238 |
| Longest exon (bp) | 20,574 |
| Mean intron in CDS length (bp) | 407 |
| Longest intron in CDS (bp) | 9,263 |
| **Protein functional annotation statistics** | |
| eggNOG | 22,214 |
| InterProScan | 22,007 |
| Pfam | 18,779 |
| GO terms | 16,423 |
| COG | 20,368 |
| Phobius | 7,053 |
| SingalP | 3,147 |
| BUSCO | 991 |
| MEROPS | 1,157 |
| CAZYme | 445 |

**Table S3. Software used for the assembly and annotation of the *Exaiptasia diaphana* (Aiptasia) strain F003 genome.**

| Software tool | Version | Source |
|---|---:|---|
| **Nuclear and mitochondrial genome assembly and evaluation** | | |
| NanoPlot | 1.42 | https://github.com/wdecoster/NanoPlot |
| Porechop | 0.2.4 | https://github.com/rrwick/Porechop |
| Flye | 2.9.3 | https://github.com/mikolmogorov/Flye |
| gfastats | 1.3.6 | https://github.com/vgl-hub/gfastats |
| Bandage | 0.9.0 | https://github.com/rrwick/Bandage |
| BlobToolKit | 4.1.2 | https://github.com/blobtoolkit/blobtoolkit |
| BLASTn / BLASTp | 2.14.1 | https://ftp.ncbi.nlm.nih.gov/blast/executables/LATEST |
| DIAMOND | 2.1.8 | https://github.com/bbuchfink/diamond |
| BUSCO | 5.8.2 | https://gitlab.com/ezlab/busco |
| GetOrganelle | 1.7.7.1 | https://github.com/Kinggerm/GetOrganelle |
| MITOS | 2.1.0 | https://gitlab.com/Bernt/MITOS/ |
| **DNA repeat identification** | | |
| RepeatModeler | 2.0.4 | https://github.com/Dfam-consortium/TETools |
| Extensive de novo TE Annotator (EDTA) | 2.2.0 | https://quay.io/repository/biocontainers/edta?tab=tags |
| RepeatMasker | 4.1.6 | https://github.com/Dfam-consortium/TETools |
| **Gene prediction and functional annotation** | | |
| funannotate | 1.8.15 | https://github.com/nextgenusfs/funannotate |
| BRAKER | 3.0.8 | https://github.com/Gaius-Augustus/BRAKER |
| tRNAscan-SE | 2.0.12 | https://github.com/UCSC-LoweLab/tRNAscan-SE |
| STAR | 2.7.11b | https://github.com/alexdobin/STAR |
| samtools | 1.16.1 | https://github.com/samtools/samtools |
| StringTie | 2.2.1 | https://github.com/gpertea/stringtie |
| InterProScan | 5.72-103.0 | https://www.ebi.ac.uk/interpro/about/interproscan/ |
| eggNOG-mapper | 2.1.12 | https://github.com/eggnogdb/eggnog-mapper |
| Phobius | 1.01 | https://phobius.sbc.su.se/data.html |
| SingalP | 6.0 | https://services.healthtech.dtu.dk/services/SignalP-6.0/ |
| AGAT | 1.4.1 | https://agat.readthedocs.io/en/latest/index.html |
| GenomeTools | 1.6.5 | https://github.com/genometools/genometools |
| GffRead | 0.12.7 | https://github.com/gpertea/gffread |
| GeneForge | 1.0 | https://github.com/SequAna-Ukon/GeneForge |
| OrthoVenn3 | 1.0 | https://orthovenn3.bioinfotoolkits.net/ |
